# Quantifying the evolutionary potential for Delta Smelt persistence in a warming habitat

**DOI:** 10.64898/2026.06.16.732742

**Authors:** Joanna S. Griffiths, Amanda J. Finger, M. Moshiur Rahman, Brittany E. Davis, Tien-Chieh Hung, Nann A. Fangue, Andrew Whitehead

## Abstract

Long-term persistence of managed species will depend, in part, on whether the species harbors the physiological or genetic potential to adjust to warming temperatures, and whether relevant genetic variation is modified by management practices. The critically endangered Delta Smelt (*Hypomesus transpacificus*) is intensively managed, but little is known about the presence of genetic variation for resistance to elevated temperature, which will be important to maintain for their persistence in a rapidly warming future. Using a pedigree and whole genome sequencing data, we characterized the genetic variation and genomic architecture for CTMax (as a metric of upper thermal tolerance) across control and elevated rearing temperatures, alongside covarying traits (body size, degree of hatchery ancestry). Warmer rearing temperatures increased CTMax through acclimation but also resulted in reduced additive genetic variation for the trait, which could constrain adaptation under thermal stress. We found that larger fish had reduced CTMax, although this effect was diminished at elevated temperatures. We observed modest heritability for CTMax at rearing temperatures of 15°C and 18°C (0.26 and 0.16, respectively), but only a limited number of loci were identified that had consistent effects on CTMax across rearing temperatures. Instead, the genomic basis of thermal tolerance was highly dependent on rearing temperature (many loci detected with a GxE effect). The influence of domestication selection was indicated by changes in allele frequency, and divergence in upper thermal tolerance and plasticity, between low and high hatchery ancestry groups. Minimal overlap between loci associated with domestication and CTMax suggests that these traits possess separate genetic underpinnings. Knowledge of genetic variation supporting ecologically relevant physiological variation may be useful for refuge management and may inform supplementation in an ever-warming environment.

## Introduction

Climate change is exposing organisms to novel environments and causing detrimental physiological impacts. Resilience to sustained and extreme temperature events will be necessary for wild species to persist, where resilience may be achieved through physiological plasticity (acclimation ability) and evolutionary adaptation. Rapid adjustments may be achieved through plasticity (Bradshaw, 1965), but plasticity may be insufficient for maintaining fitness in warming environments, such that evolution is required. A species’ capacity to evolve depends on the presence and nature of heritable genetic variation underlying putatively adaptive traits (Lynch & Walsh, 1998). Consequently, preserving genetic diversity, especially that fraction of diversity underlying adaptive traits, is a crucial objective for the conservation and management of wild species (Holderegger et al., 2006; Mable, 2019).

Genetic and environmental influences can each contribute and interact to affect phenotypic variation, where the genetic contribution can be environmentally dependent. For example, environmental stress can increase available genetic variation (e.g., expose cryptic genetic variation) or decrease genetic variation (e.g., when species approach physiological limits) (Jenkins et al., 1997; Mcguigan et al., 2010; McGuigan & Sgrò, 2009). Therefore, it is best to examine genetic variation across multiple ecologically relevant conditions. Evolutionary adaptation will also depend on available additive genetic variation for phenotypic plasticity, where genotypes may have variable effects on phenotype across environmental conditions: genotype-by-environment interactions (GxE) (Via & Lande, 1985). Another consideration is how the pace of evolutionary change may be further shaped by genetic correlations between traits, which can either constrain or facilitate evolutionary adaptation and may further depend on environmental context (Etterson & Shaw, 2001; Lande & Arnold, 1983).

Quantifying the multi-faceted drivers of evolutionary change is particularly important for at-risk species supported by conservation hatcheries or breeding programs where the primary objective is to supplement wild populations (Bradford et al., 2025; Fraser, 2008). Populations maintained in captive environments are more susceptible to losing genetic diversity due to strong genetic drift and small effective population sizes (Fiumera et al., 2004; Hedrick, 2001). Consequently, genetic bottlenecks may constrain evolutionary potential once fish are released back into the wild (Laikre et al., 2010; Lorenzen et al., 2012). These risks may be prevented or managed by genetically informed breeding that reduces the influence of genetic drift. However, captive environments can also impose strong selective pressures (intentionally or unintentionally), favoring traits like disease resistance, high yields, and an ability to live in high densities that can further erode genetic diversity.

Preserving genetic diversity through genetically informed breeding and gene flow with wild individuals has been a top priority in the captive-rearing program of the Delta Smelt (*Hypomesus transpacificus*), a critically endangered osmerid fish (California Natural Diversity Database (CNDDB), 2024; NatureServe, 2014) for which experimental releases of hatchery fish back into the wild are already underway (Baerwald et al., 2023; Davis et al., 2024; Hudson et al., 2026; T. Hung et al., 2019; U.S. Fish and Wildlife Service, 2020). The population decline and eventual crash in the Sacramento-San Joaquin Delta (Delta) was the consequence of several factors including rising temperatures and salinities, competition with non-native species, exposure to environmental pollution, and water management conflicts (Yanagitsuru et al., 2022). Delta Smelt have lower thermal tolerance limits compared to other native and non-native species in the Delta (Davis et al., 2019; Hung et al., 2022; Swanson et al., 2000), potentially further disadvantaging them relative to competitors (Halverson et al., 2022). Field experiments involving caged Delta Smelt have demonstrated that high mortality rates are driven by warm summer temperatures with a lethality threshold of 26°C (Davis et al., 2024). Given that current conditions in the wild already reach these critical limits, it is imperative to assess the extent of genetic variation for thermal resilience in captive Delta Smelt, which will determine the species’ capacity to adapt to a warming climate. This assessment is especially critical across fish with different hatchery ancestries, as recent evidence suggests that domestication may have already induced heritable shifts in Delta Smelt thermal physiology (Griffiths et al., 2026).

Our study aimed to investigate the genetic variation and genomic architecture of thermal tolerance in captive Delta Smelt. We also explored the influence of rearing temperature, hatchery ancestry (domestication index, or DI), and body size (fork length), on the architecture of this trait. Finally, we tested whether patterns of genetic variation between low and high DI fish were consistent with the influence of domestication selection. To achieve these goals, we established families of Delta Smelt from a range of hatchery ancestries (low, medium, and high DI), reared them at both ambient and elevated temperatures, measured the upper thermal tolerance of ca. 3,000 fish, and genotyped fish at thousands of loci genome-wide. We tested for genotype-phenotype associations for multiple traits including CTMax, and for trait associations that varied with rearing temperature (GxE), and compared these between fish with different degrees of hatchery ancestry.

## Methods

### Adult spawning and offspring rearing

Adult Delta Smelt were obtained from the refuge population at the University of California Davis Fish Conservation and Culture Laboratory (FCCL). Since its inception in 2008, FCCL uses genetically informed single pair crosses to reduce inbreeding and incorporates wild broodstock (when available) to maintain genetic variation (Frankham, 2010; Utter & Epifanio, 2002). Hatchery ancestry within this refuge population is quantified using a domestication index (DI) which represents the number of generations an individual’s genome has spent in captivity. Wild fish are assigned a DI of 0, and offspring produced from wild parents have a DI of 1 (Chase et al., 2024; Finger et al., 2018). Offspring DI is calculated as the parental mean DI plus 1. For the experiments reported here, adult Delta Smelt were spawned in 2021 using a modified North Carolina II breeding block design (Fig. S1), with parental pairs selected to have genetic relatedness below 0.02 (mean relatedness of crossed parents was 0.003). Eggs and sperm were manually expressed and fertilization occurred in 290-500-mL plastic bowls (freshwater, 16 ℃) (Lindberg et al., 2013). Across 13 experimental breeding blocks, eggs from two females were independently fertilized with sperm from five males, resulting in ten distinct fertilization events per block (Fig. S1). A total of 15 males and 6 females were crossed within the same DI category; low (DI < 7), medium (DI 7 –9), or high (DI > 10) (Fig. S2). Due to low fertilization success, an additional 5 males and 2 females were crossed within the low DI category. A separate “mixed” DI group was also generated by crossing males and females spanning low, medium, and high DI categories.

At three days post-fertilization (dpf), offspring from each cross were divided evenly between two rearing temperature treatments, with two replicate incubation columns at 15 ℃ and two at 18 ℃ (Fig. S1), housed within two parallel recirculating aquaculture systems (Tsai et al., 2022). Offspring from different crosses were further pooled together based on DI category (low, medium, high, or mixed), resulting in 16 total incubation columns (four DI categories × two temperatures × two system replicates). For each family, between 10 and 60 embryos were stocked per treatment replicate. Families that did not produce enough embryos to meet the minimum per-treatment allocation were excluded from the experiment. Hatching occurred at approximately 10 dpf, after which larvae were transferred to 38 L trays; at 40 dpf, larvae were moved into 133-L tanks and reared until 143–151 dpf to complete all experiments. Fish were maintained in freshwater, under a natural photoperiod, and fed live prey consisting of cultured rotifers (*Brachionus plicatus*) and brine shrimp nauplii (*Artemia franciscana*) (Lindberg et al., 2013). All animal handling, care, and experimental procedures were approved (UC Davis IACUC Protocol #21915).

### Thermal tolerance measurements

Upper thermal tolerance was assessed using a modified critical thermal maximum (CTMax) method that was consistent and repeatable (Beitinger et al., 2000; Griffiths et al., 2026) in larval Delta Smelt aged 71–79 dpf. This age was selected to maximize survival while maintaining sufficiently large sample sizes. CTMax was measured in 195 individuals from each replicate group (n ∼ 390 per treatment [4 DI groups x 2 rearing temperatures]; 3120 fish total). Individuals were placed separately into 30-mL opaque cups equipped with gentle air bubblers for mixing and aeration. Cups were then submerged in temperature-controlled water baths, where water temperature was increased at a constant rate of 0.3 ℃ min⁻¹ until fish reached loss of equilibrium (LOE), a standard endpoint for determining thermal tolerance limits (Beitinger et al., 2000; Davis et al., 2019; Jeffries et al., 2016; Komoroske et al., 2014). The water temperature at LOE was recorded using a calibrated immersion thermometer.

A linear mixed-effects ANOVA (R package lme4 v1.1-35.1 (Bates et al., 2014) and car v3.1-2 (Fox & Weisberg, 2019), was used to test for the fixed effects of DI (low, medium, or high), rearing temperature (15° or 18°C), fork length, and their interaction, on CTMax (the response variable), with trial number as a random effect, and tank/treatment group nested within the recirculating system as a random effect (Fig. S1). We also ran a linear mixed-effects ANOVA for fork length, with DI and rearing temperature as fixed effects and their interaction. To meet the assumptions of the ANOVA, we performed an orderNorm transformation on CTMax and fork length data (bestNormalize v1.9.1; Peterson, 2021).

### Family-level Survival

Using the pedigree from genotyped offspring (methodology described below), we were able to estimate the percent family survival based on the fraction of the starting number of fertilized embryos that survived to 71-79 dpf when we collected fish for CTMax measurements. We performed a linear mixed-effect ANOVA (R packages lme4 v1.1-35.1 and car v3.1-2), with family survival (percent alive) as the response variable, and DI and rearing temperature as fixed effects. The random effects in the model were the family ID and tanks that were nested within the recirculating system. To investigate how survival varied between the two rearing temperatures, we performed an ANOVA with rearing temperature and family ID as fixed effects, and tanks nested within the recirculating system as a random effect. Finally, we investigated whether family survival correlated with CTMax by estimating the mean CTMax per family per replicate. We performed a linear mixed-effect ANOVA with mean family CTMax as the response variable, mean family survival as a fixed effect, and tanks nested within the recirculating system as random effects. To meet the assumptions of the ANOVA, we performed a Box-Cox transformation on survival data (bestNormalize v1.9.1; Peterson, 2021).

### DNA extraction and library preparation

Fin clips (2-4mm^2^) were sampled from 87 parents following spawning, and from 3,085 of their offspring following CTMax trials. DNA extractions were performed using Agencourt Ampure XP beads as described in (Ali et al., 2016). DNA yield was assessed using a plate reader (BioTek Instruments, Winooski, VT). We normalized all samples to 2 ng/uL. We constructed 3,172 whole-genome sequencing libraries using a modified Twist 96-Plex Library Preparation protocol (Twist Bioscience, San Francisco, CA). We targeted an insert size of 400 bp and library quality was assessed using a Bioanalyzer (Agilent, Santa Clara, CA). We created 5 pools of 560-650 offspring samples per pool. Each pool was sequenced on a single lane of NovaSeq S4 PE150 (target depth 1-2x). One pool of all 87 parent samples was sequenced across 4 lanes of NovaSeq S4 PE150 (target depth 30x).

### Reference panel generation

To enable offspring genotype imputation, we generated a reference panel using the 87 high coverage parent samples. We used the snpArcher pipeline with snakemake v5.32.2 (https://snparcher.readthedocs.io/en/latest/index.html) to create a VCF file of the 87 parent samples. This pipeline performs quality control, read mapping, and variant calling. We mapped reads to the *Hypomesus transpacificus* reference genome (GCF_021917145.1) and called SNPs using GATK and applied the recommended GATK filtering (McKenna et al., 2010).

We applied additional filters to retain SNPs with a depth between 2-100x using vcftools v1.14 (Danecek et al., 2011) and biallelic SNPs using bcftools v1.14 (Danecek et al., 2021). We required SNPs to be genotyped in at least 50% of parents and to be in Hardy-Weinberg Equilibrium. Finally, we applied a minor allele frequency (MAF) cutoff of 2 % using vcftools (v1.14) based on our imputation accuracy (see “*Imputation Validation”* section below). We removed sites with imputation quality <0.065 (i.e., the info field generated by GLIMPSE). The filtered VCF file consisted of 1,300,115 SNPs. We used the program SHAPEIT5 v5.1.1 (Hofmeister et al., 2022) to phase all samples at common variants (MAF ≥ 0.1%). SHAPEIT5 was run with a recombination rate of 1 cM per Mb and an expected error rate in the phase informative reads of 0.0001 (the default setting).

### Offspring genotype imputation and pedigree generation

We used the same snpArcher pipeline to generate aligned bam files for the 3,085 offspring samples. We first estimated genotype likelihoods with bcftools v1.14 mpileup using a list of variant positions extracted from the parent reference panel. We then used the phased parent VCF as a reference panel to impute all low-coverage sequenced offspring samples based on genotype likelihoods. Following recommended guidelines with the GLIMPSE v1.1.1 software (Rubinacci et al., 2021), we first split the phased reference panel into smaller chunks (2Mb chunks with a buffer size of 200Kb). To impute genotypes, we ran GLIMPSE_phase with 15 iterations and then ligated chunks together using GLIMPSE_ligate. Finally, we generated a pedigree using genotyped parents and imputed offspring in the program AlphaAssign with a parentage assignment threshold corresponding to a 99.99% posterior probability (Whalen et al., 2018).

### Imputation validation

To validate imputation accuracy, we down-sampled the mapped reads from one of our parent individuals from 30x to 1x coverage using samtools v1.14 (Danecek et al., 2021) and imputed this individual using the phased reference panel. We first re-created our phased reference panel after removing this individual from the VCF file and followed the same methods as above for phasing. Next, we computed genotype likelihoods using bcftools mpileup v1.1.4 and imputed genotypes using GLIMPSE_phase for this down-sampled parent as we did for the low-coverage offspring samples. We computed the r2 correlation between imputed dosages (in MAF bins) and highly confident genotype calls. We used MAF bins of 0.00000, 0.00100, 0.00200, 0.00500, 0.01000, 0.05000, 0.10000, 0.20000, 0.50000. We were able to accurately impute variants up to ∼2 % MAF (r^2^ = 0.8 - 0.85; Fig. S3), resulting in a MAF cut-off of 2% in the phased reference panel VCF for a total of 1,300,115 SNPs.

### Quantitative Genetic Estimates

To quantify standing additive genetic variation and the capacity to evolve to warming temperatures, we estimated variance components for CTMax and fork length size using an animal model; a mixed model that uses a pedigree to estimate the relative contributions of genetic and environmental components on phenotypic variation (Kruuk, 2004). Specifically, we used a generalized linear mixed model (GLMM) using Markov chain Monte Carlo (MCMC) in the R package MCMCglmm (Hadfield, 2010) in R v.4.5.0. Using the pedigree generated from the genotyped parents and imputed offspring, we tested different model parameters to estimate variance components using DIC score comparisons for random effects (rearing temperature, dam ID, and fork length). In animal models, potential confounders (e.g., experimental system) are accounted for by including them as fixed effects and were included in the model if the 95% Highest Posterior Density (HPD) intervals for the posterior distribution for the fixed variables did not overlap zero. Our final model for estimating variance components for CTMax and fork length included experimental system as a fixed effect and rearing temperature as a random effect. Rearing temperature was set as a covariate random effect in the model so we could further partition the additive genetic variation by rearing temperature as a random interaction so that each animal has a separate effect for each rearing temperature level. This allowed us to directly compare variance components and heritability across different rearing temperatures in a single model. We ran these models with all fish, as well as separate identical models with the data subset by each DI category (low, medium, and high). Given that maternal effects have been shown to influence size of offspring in other species (Moore et al., 2019; Paul et al., 2023), we ran an additional model for fork length with dam ID as an additional random effect even though it was not statistically supported in model selection criteria.

To assess the extent to which additive genetic effects underlying thermal tolerance were conserved across rearing temperatures, we calculated additive co-variation and genetic correlation for family-level mean CTMax measured at 15℃ and 18℃. We fit a multivariate model using full-sibling family mean CTMax measured at 15 ℃ and 18 ℃ as co-varying traits. In this model, tank replicate was included as a fixed effect. This approach enabled us to directly estimate the genetic correlation between family genotype and CTMax performance at 15°C vs. 18°C rearing. A correlation near 0 indicates that thermal performance at 15°C does not predict performance at 18°C, which suggests that there is some re-ranking of genotype performance at each rearing temperature.

To quantify genetic correlations among traits and estimate additive genetic co-variation, we fit two additional multivariate animal models. Phenotypic correlations arise from multiple sources, including shared environments, maternal effects, and additive genetic covariance, whereas genetic correlations specifically reflect shared additive genetic influences between traits, arising from pleiotropy or physical linkage among loci. Therefore, we fit a multivariate model with CTMax and fork length as the co-varying response traits, and a second model with CTMax and numerical DI as the co-varying response traits. In both models, experimental system was included as a fixed effect and rearing temperature as a random effect. These analyses allowed us to assess whether genetic variation underlying thermal tolerance covaries with body size or DI across the two rearing temperatures.

All models were run for 500,000 iterations, a burn-in of 1,000, and a thinning interval of 500, resulting in a posterior sample size of 998 estimates. Autocorrelation values for the parameters were near zero, confirming that convergence occurred and there were no trends in the parameters over successive generations of the model. Heritability was calculated as the ratio of observed additive genetic variance to total phenotypic variance. We also calculated evolvability for CTMax and fork length as the ratio of observed additive genetic variance to the squared mean of the trait value. Genetic correlation between two traits was calculated as the ratio of observed additive genetic co-variation to the square root of additive genetic variation for the two traits multiplied together. All quantitative genetic estimates are reported as posterior modes with the 95% HPD intervals.

### Genomic architecture of thermal tolerance and fork length

We performed a genome-wide association study (GWAS) for thermal tolerance and fork length using the program GEMMA v0.98.3 (Zhou & Stephens, 2012). A relatedness matrix was computed in GEMMA with the centered method. To test for the main effect of genotypes on CTMax and fork length, we ran a linear mixed model with the trait measurement as the response variable and rearing temperature and experimental system as covariates. We were unable to include fork length as an additional covariate in the model given that it is highly correlated with the temperature treatments in the experimental system covariate parameter (i.e., growth was faster, and therefore size was larger, in fish reared at elevated temperature). GEMMA also reports the proportion of variance explained by genotypes (“SNP heritability”) for the trait from the linear mixed model. To detect genotypic effects on CTMax that depended on the rearing temperature (GxE), we ran a second linear mixed model with the “-gxe” flag. The linear model controls for both the SNP main effect and environmental (rearing temperature) main effect, while testing for the interaction (i.e., genetic effect on CTMax or fork length that is temperature-dependent). We plotted the results of the linear models as a manhattan plot (R package qqman) and applied a genome-wide and chromosome-level Bonferroni correction.

### Signatures of domestication selection

We investigated evidence for genomic divergence between low and high domesticated fish using F_ST_ and GWAS approaches. We measured pairwise F_ST_ (Weir methodology, window size of 10,000 bp, sliding window of 5,000 bp) between low and high DI progenitor fish (N=31 and 29, respectively) using vcftools v0.1.16 (Danecek et al., 2011). We ran a permutation test to determine whether F_ST_ was higher than expected by chance by randomly assigning individuals into two groups (1,000 random permutations) as in Griffiths et al. (2026).

We also explored genomic associations with hatchery ancestry in the offspring dataset using a GWAS on numerical DI with methods similar to those described above for GWAS of thermal tolerance and fork length. We ran a linear mixed model in GEMMA with the numerical DI as the response variable and experimental system as a covariate. We plotted the results of the linear models as a manhattan plot (R package qqman) and applied a genome-wide and chromosome-level Bonferroni correction.

### Functional annotation and enrichment of candidate SNPs

We used the program LD-annot v0.4 (Prunier et al., 2019) with r^2^ = 0.8 to identify gene regions and their annotations that are in linkage disequilibrium (LD) with SNPs statistically associated with CTMax, fork length, and DI GWAS analyses, and the F_ST_ analysis. Gene Ontology (GO) terms for each gene were identified by matching Delta Smelt gene names with those from the zebrafish annotation database in ensembl using the R package biomaRt v2.46.3 (Durinck et al., 2005, 2009). GO enrichment analysis was performed using the R package topGO v2.42.0 (Alexa & Rahnenführer, 2007), with the background gene set defined as genes in LD with SNPs from the filtered progenitor VCF (23,835 genes). We used a Fisher’s Exact test (p<0.05) and retained GO terms that were represented by more than one significant gene.

To further explore the potential functions and downstream effects of genes associated with significant SNPs in the GWAS and F_ST_, we investigated whether these genes overlapped with those that were altered in their transcription or methylation by rearing temperature or DI; in a companion study, we had characterized differentially expressed genes (DEGs) and differentially methylated regions (DMRs) between 15℃ and 18℃ rearing temperatures and between low and high DI fish (Griffiths et al., 2026). For candidate genes identified in our GWAS for CTMax, we compared overlap with DEGs and DMRs due to rearing temperature. For candidate genes identified in our offspring GWAS for DI and our progenitor F_ST_, we compared overlap with DEGs and DMRs due to DI. We used a hypergeometric test (phyper function in base R v4.3.2) to determine whether the significant overlap was more than expected by chance.

### Shared genomic basis

We evaluated whether traits shared a common genomic basis. We first evaluated overlap among sets of genes significantly associated with phenotypes in GWAS analyses, including the genetic-only associations and temperature-dependent associations for both CTMax and fork length, and the genetic-only association with DI. To standardize among different statistical thresholds used in each GWAS, we selected the top 300 SNPs that were associated with each trait and identified genes in LD with those SNPs using LD-annot v0.4 as described above. We also compared the top 300 SNPs from each GWAS to the low-high progenitor F_ST_. We used two F_ST_ thresholds for comparisons; the top F_ST_ windows containing 300 SNPs and SNPs within windows with an F_ST_ above 0.1. We then used a hypergeometric test (implemented with the *phyper* function in base R v4.3.2) to assess whether the observed gene overlap exceeded expectations under random association, given the total number of genes (23,835) in LD to SNPs in the VCF. In addition, we applied a more stringent approach at the SNP level using the program PLACO+ (Park & Ray, 2026). This approach also accounts for correlation among traits that arises from measurements on the same individuals. PLACO+ uses GWAS summary statistics and tests each SNP for joint association with both traits, returning a p-value that indicates whether a SNP is significantly associated with both traits. We applied genome-wide and chromosome-level Bonferroni correction.

## Results

### Family survival, CTMax, and fork length

A total of 130 families were generated, but 7 families had 0% fertilization success, and 14 families had poor fertilization success (defined by fewer than 40 total fertilized embryos). Out of the 3085 individuals genotyped, there were 104 families represented. Survival rates among families were similar across rearing temperatures (F_1,10.3_ = 0.92 p = 0.36; Fig. S4A) and across DI categories (F_1,57_ = 0.67 p = 0.68; Fig. S4A). We also investigated the plasticity of family survival under different rearing temperatures. However, percent survival rates were significantly variable among different families (F_1,102_ = 5.8 p < 2e-16; Fig. S4B) and there was significant variation in resilience to temperature among families (family-level survival by rearing temperature interaction; F_1,94_ = 1.6 p = 0.005; Fig. S4B).

Rearing fish at 18 ℃ compared to 15 ℃ significantly increased mean CTMax values by 0.6℃ (F_1,17.4_ = 26.2 p = 7.9e-5; Fig. 1A) and higher DI fish had higher CTMax values (F_1,42.2_ = 23.9 p = 1.5e-5; Fig. 1A), with a 0.24℃ increase in CTMax with each 1-unit increase of DI (∼1 generation spent in the hatchery). However, there was no interaction effect of rearing temperature and DI (F_1,55.4_ = 0.13 p = 0.71; Fig. 1A). Fork length was also significantly correlated with CTMax values (F_1,2109_ = 119.6 p = 2.2e-16; Fig. 1B), with a significant interaction of rearing temperature and fork length (F_1,1682_ = 131.6 p = 2.2e-16; Fig. 1B); larger fish had a smaller CTMax value, but this effect was only present when fish were reared at 15 ℃. When investigating family-level trait means, families with higher percent survival were also more likely to have higher mean CTMax (F_1,384_ = 22.4 p = 3.1e-06; Fig. S5).

**Figure 1:**
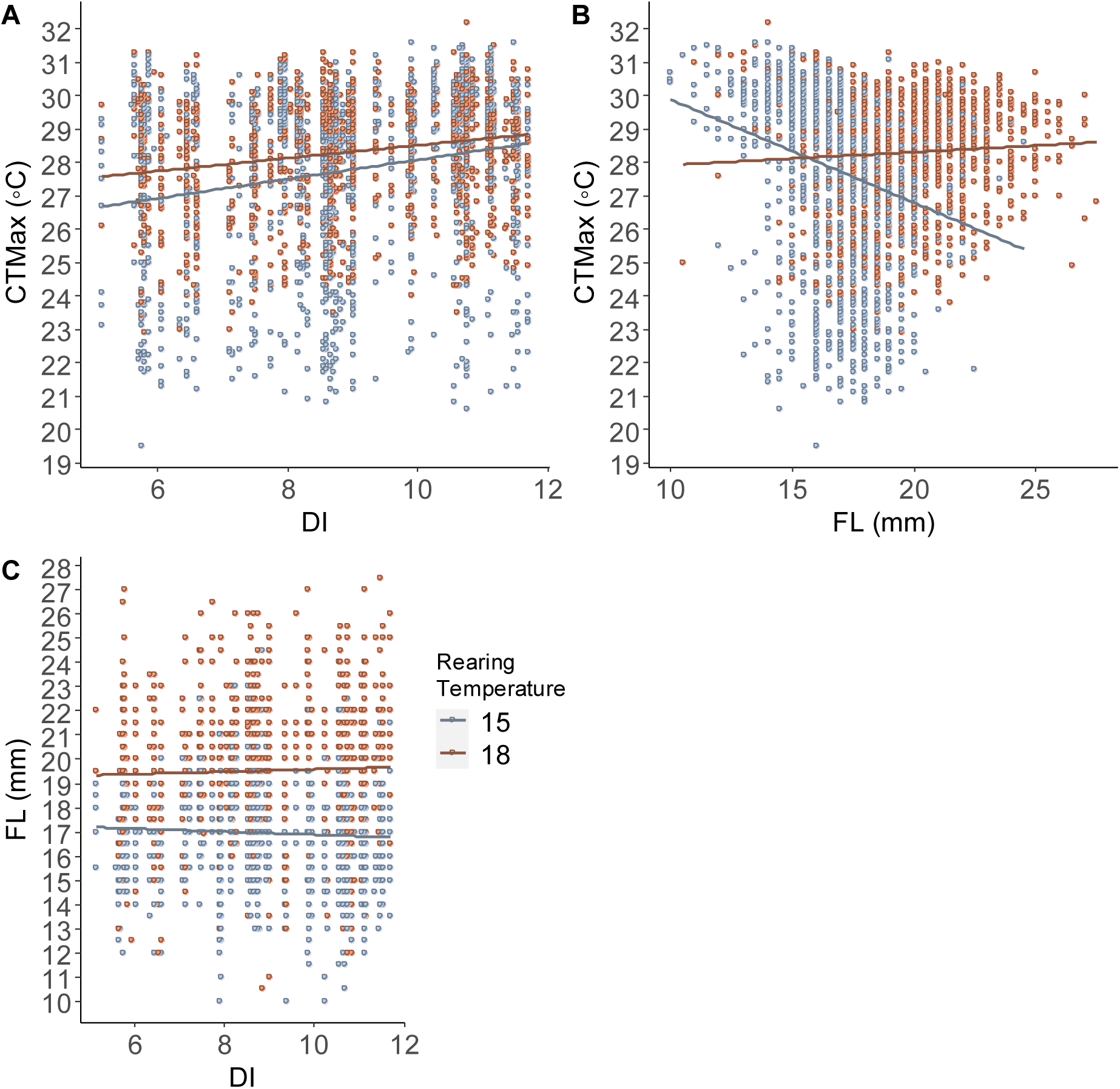
Critical Thermal Maximum (CTMax) for fish reared at either 15°C (blue) or 18°C (orange). A) Fish reared at 18°C had a higher CTMax than fish reared at 15°C and high domestication index (DI) fish had higher CTMax than low DI fish. B) Negative correlation was detected between CTMax and size (fork length, FL), where larger fish had lower CTMax. However, this correlation only existed for fish reared at 15°C. C) Fish reared at 18° C were larger than fish reared at 15°C, but there was no correlation between fork length and DI.

Rearing fish at 18 ℃ compared to 15 ℃ significantly increased mean fork length by 2.6 mm (F_1,14.1_ = 17.5 p = 9.0e-4; Fig. 1C). DI had a statistically significant effect on fork length (F_1,1257_ = 49.2 p = 3.66e-12; Fig. 1C), but the slope is close to zero (0.15mm of increase in FL per 1 unit change in DI), so we infer a very minor biological effect. Finally, there was no interaction effect of rearing temperature and DI on fork length (F_1,1119_= 2.58 p = 0.23; Fig. 1C).

### Quantitative genetics estimates

We calculated the amount of additive genetic variation, heritability (h^2^), and evolvability for CTMax and fork length in Delta Smelt progeny. Additive genetic variation (V_A_) is the component of phenotypic variation that is attributable to the additive linear effects of individual alleles and represents the genetic component that responds to natural selection, while heritability expresses V_A_ as a proportion of phenotypic variation and is used to explain how much of the observed differences in a trait is genetically determined. Evolvability represents the capacity of a trait to evolve relative to the population’s mean. Unlike heritability, which is influenced by environmental variance, evolvability is the mean-scaled measurement of V_A_, providing a more direct representation of evolutionary potential (Hansen et al., 2011).

The additive genetic variation for thermal tolerance (CTMax) for fish reared at 15℃ was significantly higher (1.91, 95% HPD: 1.02 – 2.70; Fig. 2A) than for fish reared at 18℃ (0.53, 95% HPD: 0.29 - 0.87; Fig. 2A). Similarly, the estimates of evolvability for fish reared at 15℃ was significantly higher (0.25 %, 95% HPD: 0.14 – 0.36 %; Fig. 2C) than for fish reared at 18℃ (0.065 %, 95% HPD: 0.04 - 0.1 %; Fig. 2C). However, our estimates of heritability were not significantly different between fish reared at 15℃ (0.26, 95% HPD: 0.17 - 0.40) or 18℃ (0.16, 95% HPD: 0.11 - 0.30; Fig. 2B). There was no significant difference in V_A_ for fork length between fish reared at 15℃ (2.22, 95% HPD: 1.47 – 3.34; Fig. 2D) or 18℃ (3.48, 95% HPD: 1.99 - 5.16; Fig. 2D). Similarly, there was no significant difference in heritability estimates for fork length between fish reared at 15℃ (0.78, 95% HPD: 0.50 – 1.16; Fig. 2E) or 18℃ (0.74, 95% HPD: 0.57 - 1.42; Fig. 2E), nor were there significant differences in evolvability for fork length in fish reared at 15℃ (0.25 %, 95% HPD: 0.14 – 0.36 %; Fig. 2F) and at 18℃ (0.065 %, 95% HPD: 0.04 - 0.1 %; Fig. 2F). When dam ID was included as a random effect in the model, posterior modes were lower but not significantly different than the model without dam effects (Fig. 2B). For both CTMax and fork length traits, when the data was subset for each DI category, there were no significant differences in additive genetic variation, heritabilities, nor evolvability between fish reared at 15℃ or 18℃, nor among different DI categories (Fig. 2A, B). However, we note that the HPD intervals were large, suggesting that statistical power for detecting DI effects was low.

**Figure 2.**
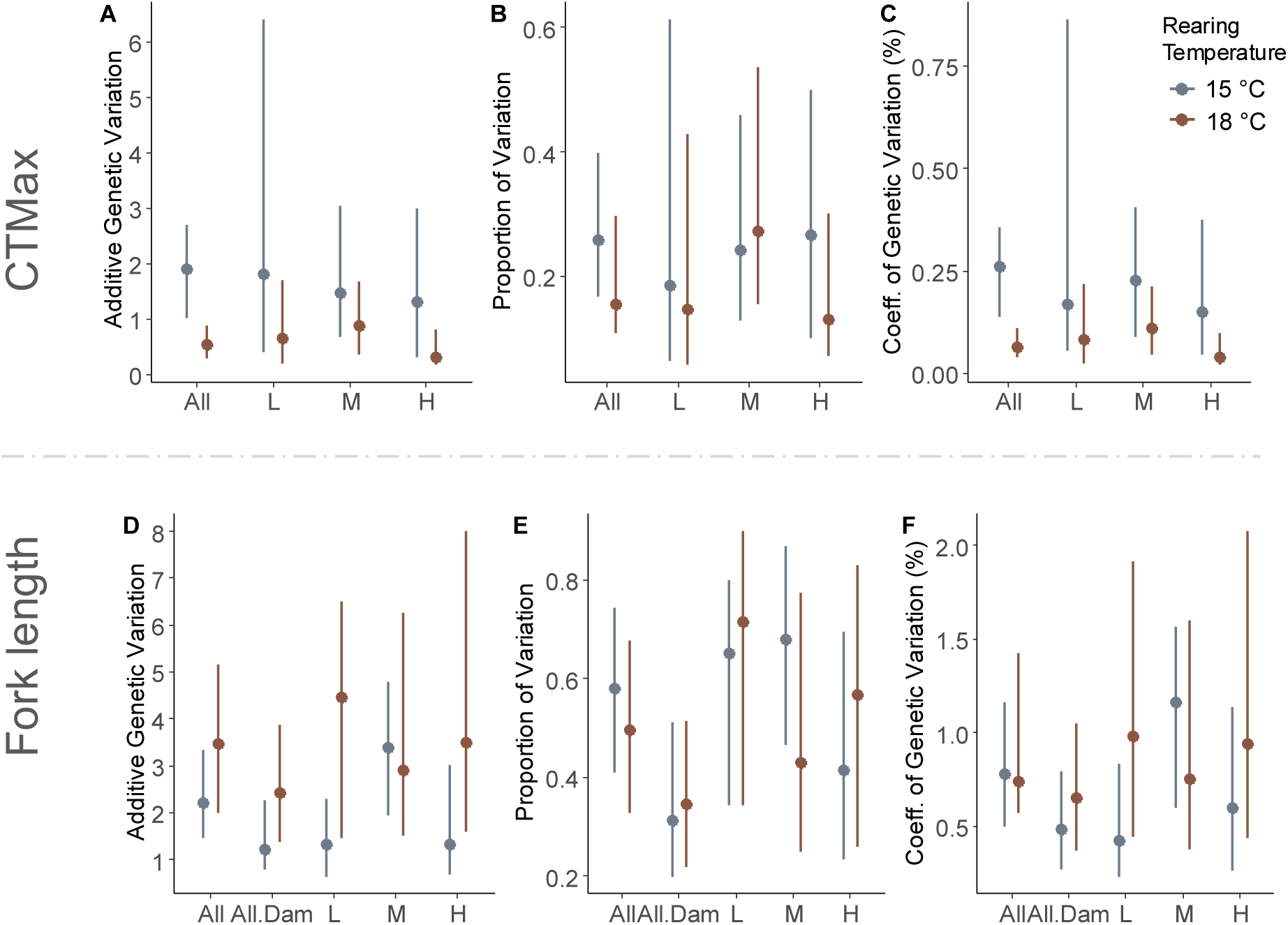
Additive genetic variation for CTMax (A) and fork length (D), proportion of phenotypic variation attributed to additive genetic variation (narrow-sense heritability; h^2^) for CTMax (B) and fork length (E), and evolvability (coefficient of genetic variation, %) for CTMax (C) and fork length (F). Points represent the mode of posterior distributions from the animal model with 95% confidence intervals displayed. Results are presented for the two models run with all fish (All: without dam effects; All.Dam: with dam effects), and data subset by each domestication index (DI) category (L: low, M: medium, and H: high). A model with all fish and Dam ID set as a random effect for fork length are shown in (D, E, and F). Variation was further partitioned by fish reared at 15℃ (blue) or 18℃ (orange).

To assess the extent to which additive genetic effects underlying thermal tolerance were conserved across rearing temperatures, we calculated additive co-variation and genetic correlation for family-level mean CTMax measured at 15℃ and 18℃. The additive co-variation between family-level CTMax at the two temperatures was significantly positive (0.30; 95% HPD 0.05 - 0.50) and the estimated genetic correlation was also positive (0.46; 95% HPD 0.08 - 0.73). However, we note that these two traits are not perfectly correlated indicating that the genetic ranking of thermal tolerance also changes in each rearing temperature (i.e., genotype-by-environment interaction).

We also calculated the genetic correlation between thermal tolerance and fork length and numerical DI (the shared genetic basis between two separate traits). We found a strong negative genetic correlation between CTMax and fork length when fish were reared at 15℃ (-0.50, 95% HPD: -0.69 to -0.24), but no genetic correlation when fish were reared at 18℃ (-0.04, 95% HPD: -0.32 – 0.28). However, we note that the HPD overlap slightly, suggesting that there is not a statistically significant difference in genetic correlations at each rearing temperature. We found a slightly positive genetic correlation between CTMax and numerical DI when fish were reared at 15℃ (0.07, 95% HPD: 0.02 - 0.16) and 18℃ (0.1, 95% HPD: 0.02 – 0.17). However, there was no difference in genetic correlations for CTMax and DI between the 15℃ and 18℃ rearing temperatures.

### Genomic architecture of thermal tolerance

A GWAS was used to identify the loci underlying variation in thermal tolerance. We tested for the effect of genotypic variation on CTMax (while controlling for rearing temperature) and genotypic effects on CTMax that depended on rearing temperature (GxE). The phenotypic variation explained by genotype (i.e., “SNP heritability”) was estimated at 0.14 (SE ± 0.03). In the GWAS there was a single SNP on an unplaced scaffold that passed the significance threshold (Fig. 3). This SNP was in LD with two protein coding genes, the uncharacterized protein unc13bb and the chemokine protein ccl19b. The genomic architecture of thermal tolerance has been widely studied in many species and suggests that it is largely polygenic (Fuller et al., 2020; Gonen et al., 2024; Griffiths et al., 2020; Michalak et al., 2019; Tobler et al., 2014). Therefore, we hypothesize that there are likely many more loci associated with thermal tolerance in Delta Smelt that did not pass the significance thresholds but that were still distinguished as peaks of association compared to the rest of the genome. Accordingly, we performed a functional enrichment test on the genes linked to SNPs that passed the chromosome-level significance threshold (62 SNPs that are in LD with 26 genes), as well as genes linked to SNPs in the top 0.04% of associations (547 SNPs that are in LD with 209 genes). This threshold was associated with a -log p-value of 3.5 (red line in Fig. 3) and was chosen to encapsulate peaks of association (“skyscrapers”) of many SNPs. Genes linked with SNPs that passed the chromosome-level significance threshold were not enriched for any GO terms. Additionally, SNPs on chromosomes 24 and 25 were either not in linkage with any genes or were unannotated. Loci passing the -log p-value of 3.5 threshold were enriched for Biological Processes GO terms including *chromatin remodeling and organization*, and *protein-DNA complex organization*, and for Cellular Components *golgi apparatus, (intracellular) membrane-bounded organelle, (intracellular) non-membrane-bounded organelle, intracellular organelle, nuclear protein-containing complex, catalytic complex, and transferase complex* (Table S1).

**Figure 3:**
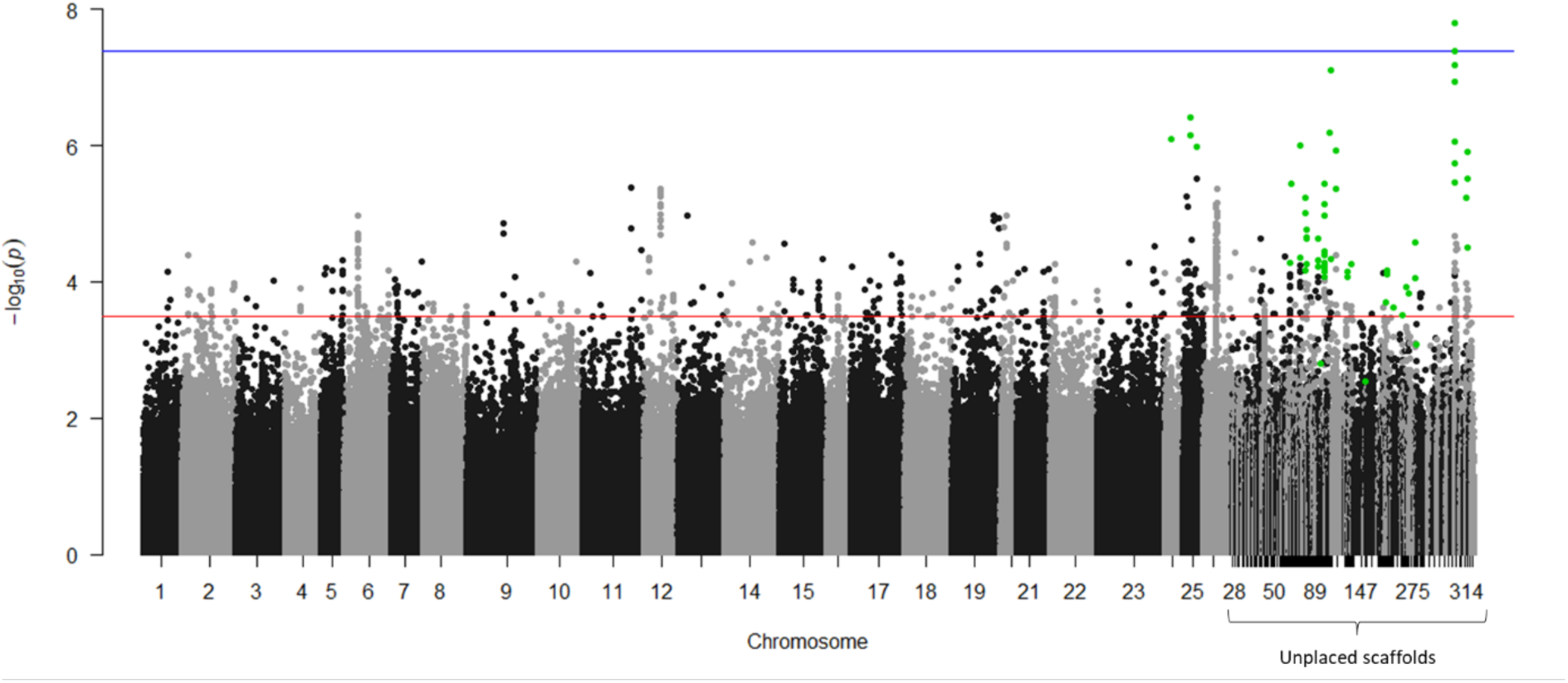
Genome-wide association test (GWAS) for the effect of genotypic variation on CTMax scores for 3,085 individual fish. Manhattan plot of *GEMMA* model results (-log pvalue) for SNPs associated with CTMax. The blue horizontal line represents the genome-wide p=0.05 significance threshold after Bonferroni correction. The red horizontal line at -log p-value 3.5 was chosen to encapsulate peaks of association for functional enrichment exploration. SNPs highlighted green are ones that pass the chromosome (or scaffold) p<0.05 Bonferroni corrected significance threshold. Genes linked to SNPs highlighted in green and those above the red horizontal line were tested for GO enrichment.

For the temperature-dependent genetic effects (GxE) on CTMax GWAS, we identified 8 independent loci across 7 chromosomes that passed the genome-wide p-value threshold (Fig. 4). These loci overlapped three gene regions, elongation factor Tu mitochondrial (tufm), reticulophagy regulator 3 (retreg3), and cell cycle progression protein 1 (ccpg1). We tested for functional enrichment of genes linked to SNPs that passed the chromosome-level significance threshold (314 SNPs, excluding SNPs found on contigs, that are in LD with 150 genes). These genes were enriched for 12 Biological Processes GO terms including *(transmembrane) transport, translation, peptide metabolic process, monoatomic ion (transmembrane) transport, peptide biosynthetic process, amide metabolic process, amide biosynthetic process,* and *(establishment of) localization* (Table S2). There were 15 Molecular Function GO terms enriched, including *GTPase activator/regulator activity, (ion transmembrane) transporter activity, enzyme activator/regulator activity, channel, catalytic activity acting on RNA*, and *salt transmembrane transporter activity* (Table S2).

**Figure 4:**
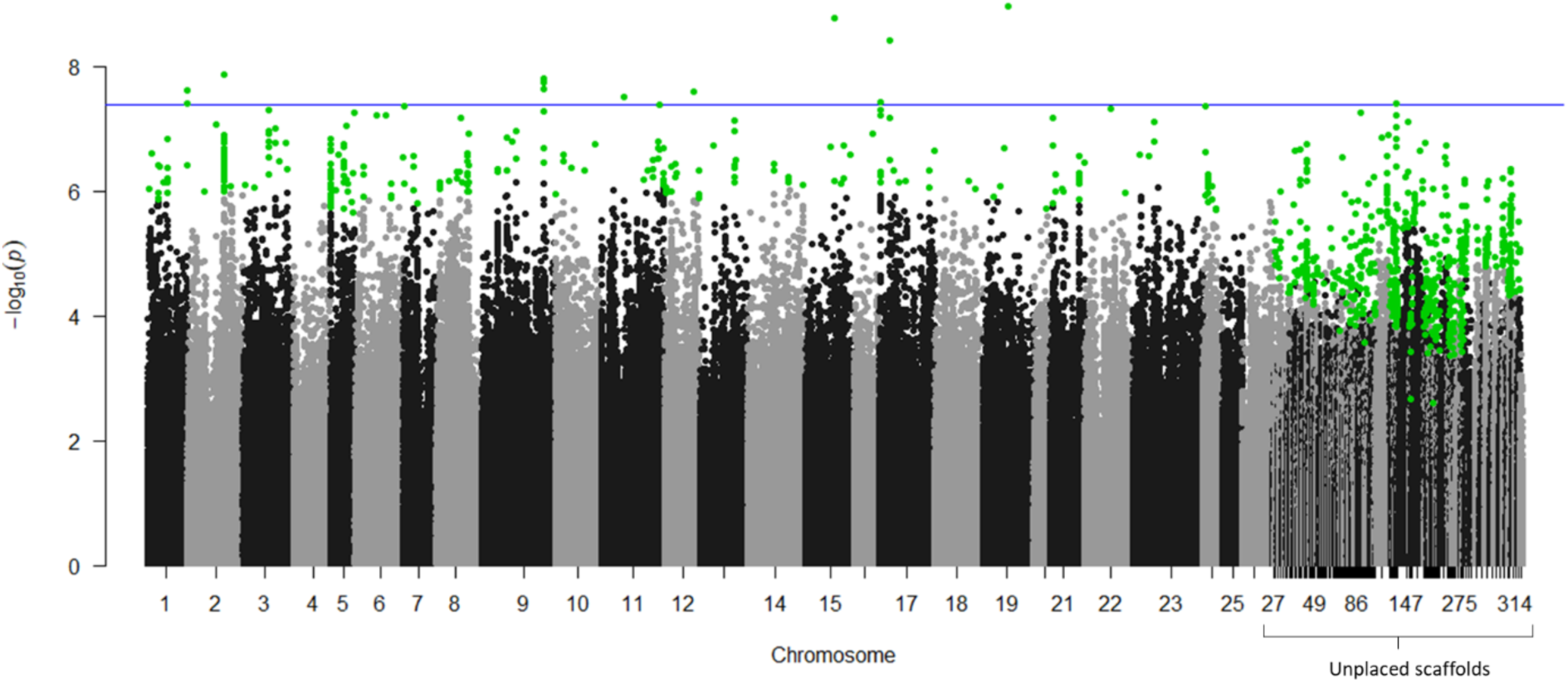
Genome-wide association test (GWAS) for temperature-dependent genetic effect on CTMax scores. Manhattan plot of *GEMMA* model results (-log pvalue) for GxE interactions associated with CTMax. The blue horizontal line represents the genome-wide p=0.05 significance threshold after Bonferroni correction. SNPs highlighted green are ones that pass the chromosome (or scaffold) p<0.05 Bonferroni corrected significance threshold.

### Genomic architecture of fork length

A GWAS was used to reveal loci underlying variation in fork length. We tested for the effect of genotypic variation on fork length (while controlling for the rearing temperature) and the genotypic effects on fork length that depended on the rearing temperature (GxE). The phenotypic variation explained by genotype (i.e., “SNP heritability”) was estimated at 0.30 (SE ± 0.03). We detected one locus of large effect on chromosome 8 (Fig. 5). This locus was in LD with four protein coding genes, two of which were annotated as zinc finger C2HC domain-containing protein 1A (zc2hc1a) and two annotated as contactin associated protein 2 (cntnap2a). We did not find any other SNPs that passed genome-wide or chromosome-level significance thresholds (except for on unplaced scaffolds). However, there were many peaks of association that may contribute to variation in fork length but were of lower effect size and below statistically significant thresholds. We therefore performed a functional enrichment test on the genes linked with SNPs that passed the chromosome-level significance threshold (71 SNPs that are in LD with 15 genes), as well as those linked with SNPs in the top 0.05 % of associated loci (613 SNPs that are in LD with 192 genes). This cut-off was associated with a -log p-value of 3.5 (red line in Fig. 5) and was chosen to encapsulate stacked peaks of association (“skyscrapers”) of many SNPs. SNPs passing the chromosome-level significance threshold were not enriched for any GO terms. Genes associated with SNPs that passed the -log p-value of 3.5 threshold were enriched for Biological Processes GO terms including *organelle organization, cellular component organization, cellular homeostasis, homeostatic process, cellular lipid metabolic process, glycerolipid process,* and *regulation of biological quality.* Enriched Molecular Processes GO terms included *protein binding* and *hydrolase activity,* and enriched Cellular Component GO terms included *receptor complex, membrane protein complex, plasma membrane protein complex,* and *plasma membrane signaling receptor complex* (Table S3).

**Figure 5:**
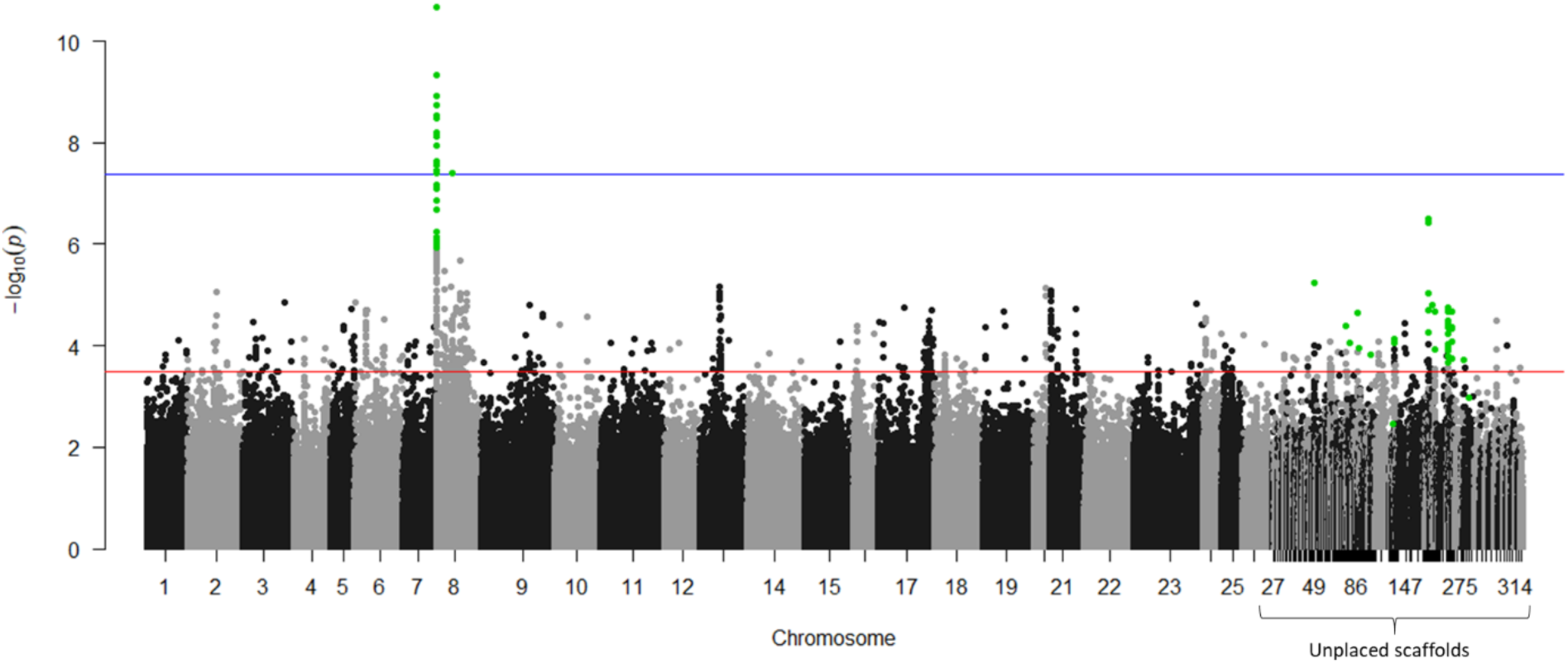
Genome-wide association test (GWAS) for the effect of genotypic variation on fork length. Manhattan plot of *GEMMA* model results (-log pvalue) for SNPs associated with fork length. The blue horizontal line represents the genome-wide p=0.05 significance threshold after Bonferroni correction. The red horizontal line at -log p-value 3.5 was chosen to encapsulate peaks of association for functional enrichment exploration. Genes linked to SNPs highlighted in green and those above the red horizontal line were tested for GO enrichment.

We identified many SNPs that had a temperature-dependent effect on fork length, suggesting that the genomic basis of body size may also depend on the rearing temperature (Fig. 6). Many SNPs passed both the genome-wide and chromosome-level threshold. We investigated the functional enrichment of SNPs that passed the genome-wide threshold only (828 SNPs in LD with 268 genes). These SNPs were functionally enriched in Biological Processes terms, including *microtubule-based movement,* and *carbohydrate metabolic process,* and Molecular Function terms, including *(phosphoric ester) hydrolase activity, phosphatase activity, actin (filament) binding, (small) GTPase binding, microtubule motor activity,* and *cytoskeletal protein binding* (Table S4).

**Figure 6:**
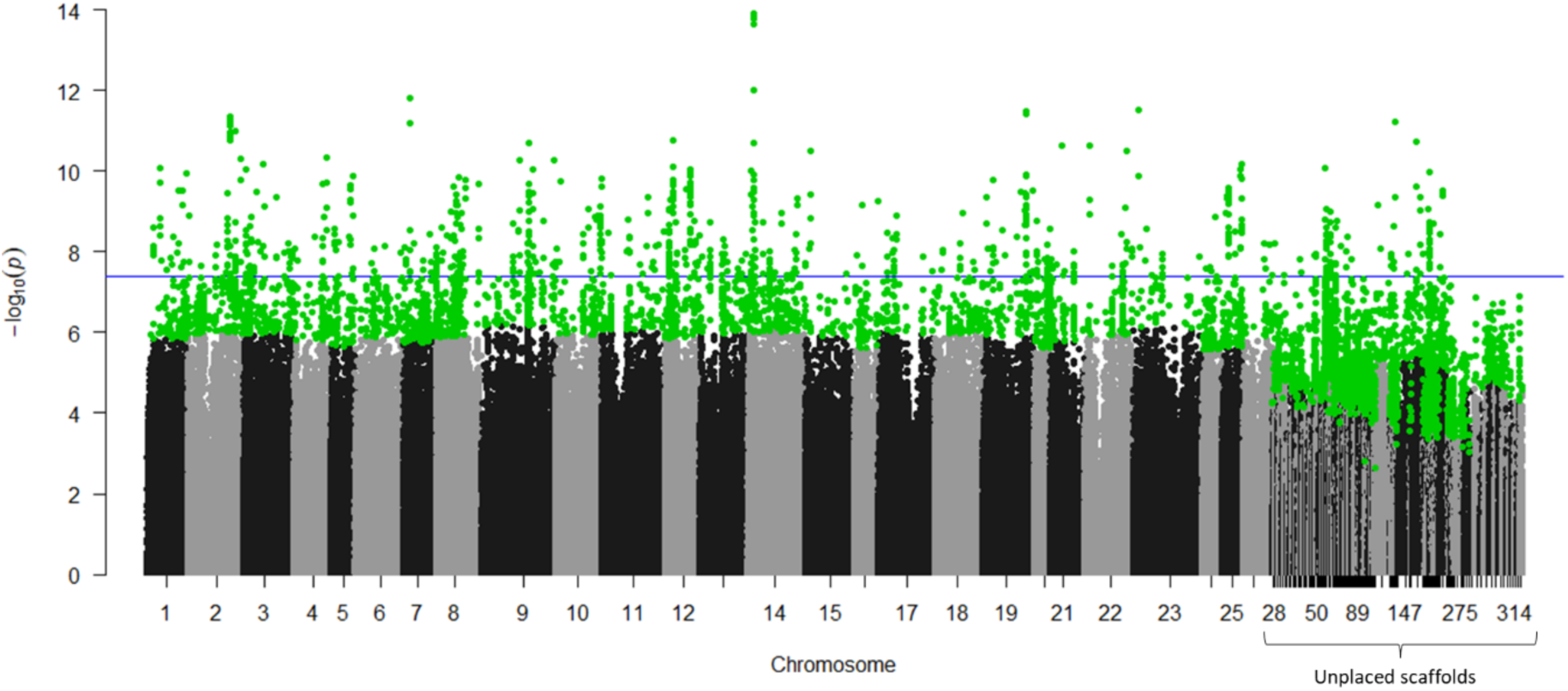
Genome-wide association test (GWAS) for genetic effects that are temperature-dependent on fork length. Manhattan plot of *GEMMA* model results (-log pvalue) for GxE interactions associated with fork length. The blue horizontal line represents the genome-wide p=0.05 significance threshold after Bonferroni correction. SNPs highlighted in green pass the chromosome-level Bonferroni correction threshold. Genes linked with SNPs above the genome-wide threshold were tested for GO enrichment.

### Signatures of domestication selection

We detected evidence of domestication selection in the hatchery, as measured by F_ST_ between low and high DI groups of progenitor fish (N=31 and 29, respectively). Genome-wide average F_ST_ was low (0.008) but statistically significantly greater than zero (p-value =0; all permutated F_ST_ values were below 0.008). Additionally, some regions of the genome across almost all chromosomes displayed peaks of F_ST_ elevated above 0.1 (Fig. 7). We performed a functional enrichment test on the SNPs in the top 0.3% of the sliding window regions (6740 SNPs that are in LD with 210 genes). This threshold was associated with F_ST_> 0.1 (Fig. 7) and was chosen to encapsulate regions of elevated divergence (“skyscrapers”). Genes linked with these high-differentiation SNPs were enriched for Biological Processes GO terms including *chemical synaptic transmission, multicellular organism, development, anatomical structure, development, cell-cell signaling, multicellular organismal process, developmental process, anterograde trans-synaptic signaling, trans-synaptic signaling*, and one Molecular Function GO term *DNA-binding transcription factor activity* (Table S5).

**Figure 7.**
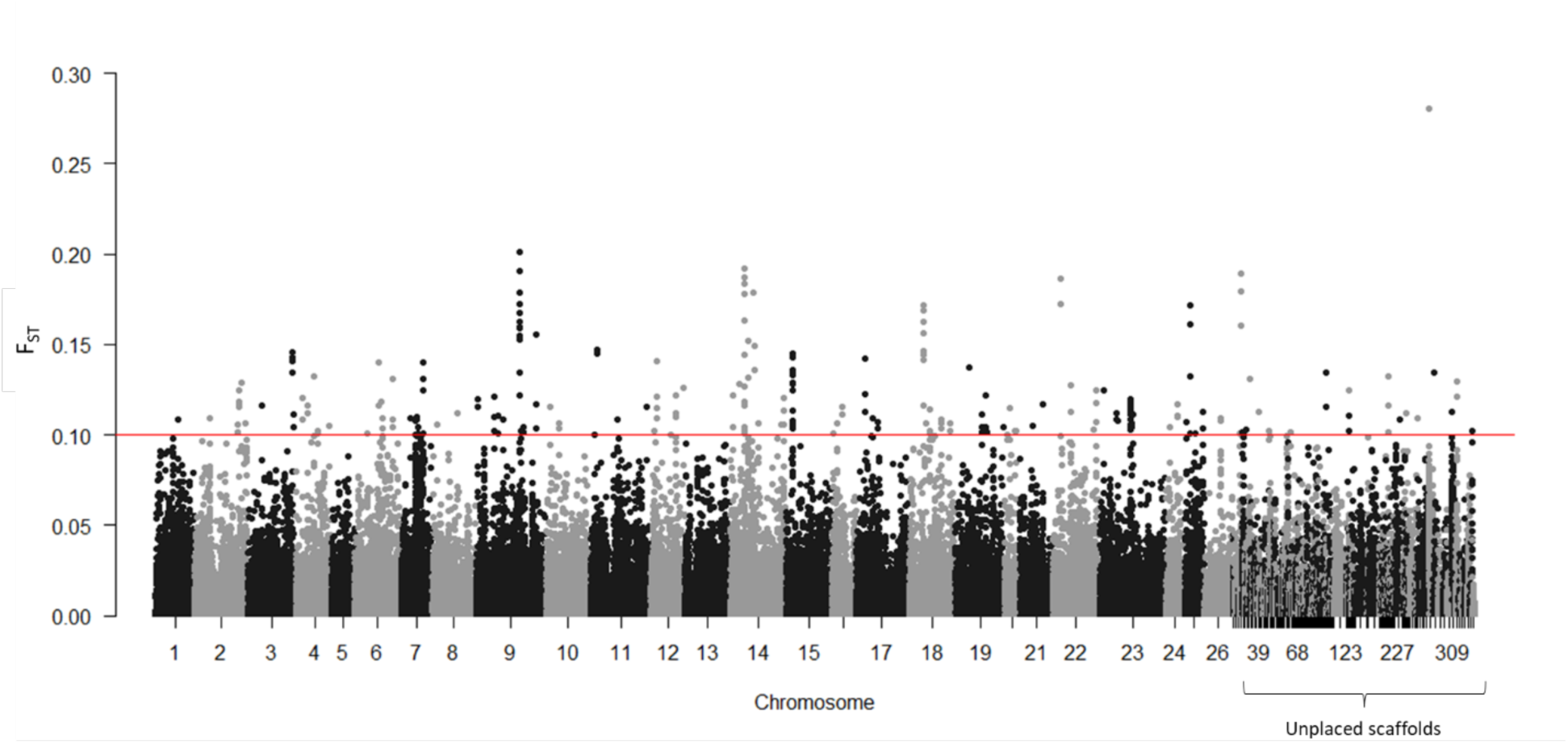
Manhattan plot of F_ST_ average across each of 10,000-bp sliding windows between low and high domestication index (DI) adult progenitors. Genes linked with SNPs within windows above the red horizontal line (F_ST_>0.1were tested for GO enrichment.

We also investigated loci associated with domestication (DI) in offspring using a GWAS approach that allowed us to account for relatedness among individuals. We identified many loci across multiple chromosomes associated with DI that passed the genome-wide and chromosome-level significance thresholds (Fig. 8). We performed a functional enrichment test on genes linked with the SNPs that passed the genome-wide significance threshold (147 SNPs that are in LD with 69 genes), as well as with loci that passed the chromosome-level significance threshold (224 SNPs, excluding contig SNPs, that are in LD with 165 genes). For those passing the genome-wide threshold, genes were enriched for the Biological Processes GO term *vesicle-mediated transport*, Molecular Function GO terms including, (*protein-macro) molecule adaptor activity,* and Cellular Component GO terms including, *plasma membrane* and *cell periphery* (Table S6). For those passing the chromosomal-level significance threshold, genes were enriched in Biological Processes GO terms, including *(homophilic) cell adhesion,* and *cell-cell adhesion.* Enrichment in Molecular Function GO terms included, *protein-macromolecule adaptor activity, phosphatidylinositol binding,* and *molecular adaptor activity.* Enrichment in Cellular Component GO terms included, *cell periphery,* and *plasma membrane protein complex* (Table S7).

**Figure 8.**
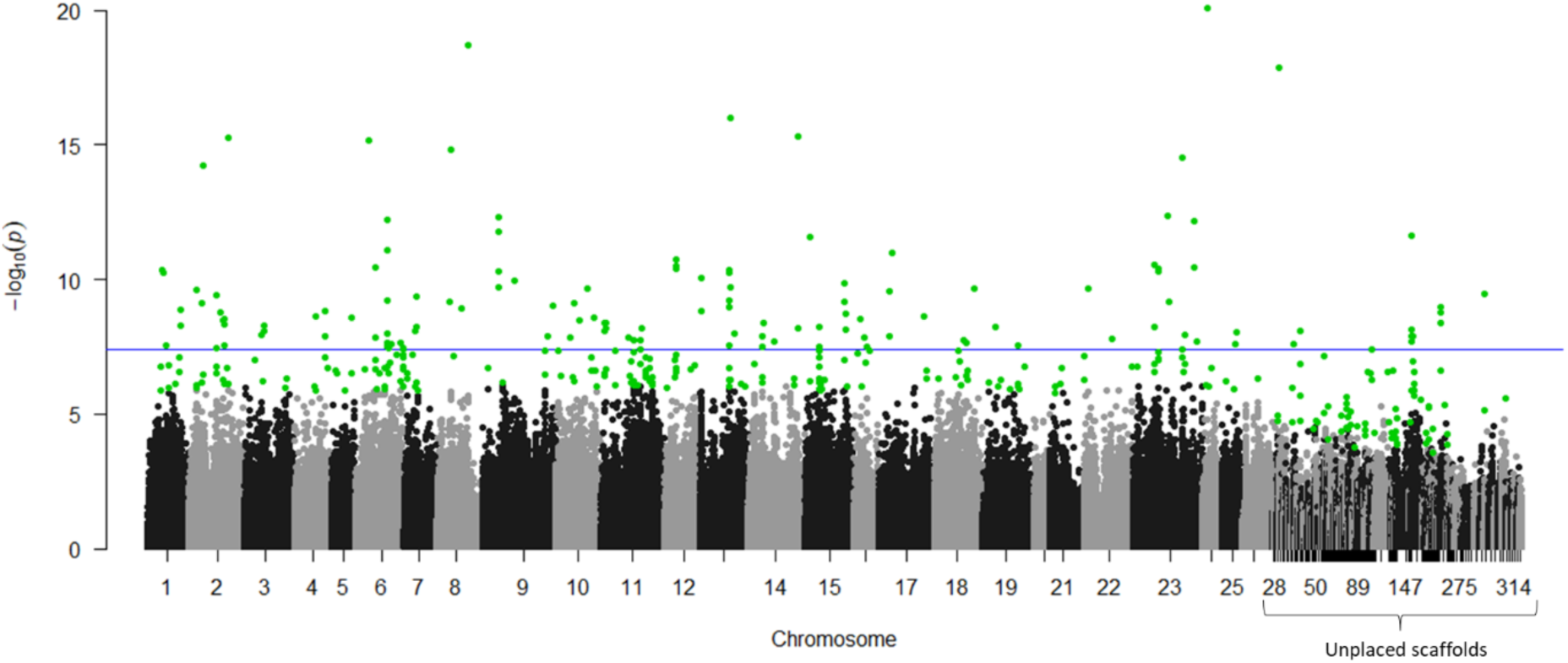
Genome-wide association test (GWAS) for the effect of genotypic variation on domestication index (DI) in offspring. Manhattan plot of *GEMMA* model results (-log pvalue) for SNPs associated with numerical DI value. The horizontal blue line represents the genome-wide p=0.05 significance threshold after Bonferroni correction. SNPs highlighted green are those that pass the chromosome (or scaffold) p>0.05 significance threshold following Bonferroni correction. Genes linked with SNPs highlighted in green and those above the blue horizontal line were tested for GO enrichment.

### Minimal shared genomic basis among traits

Pedigree-based animal models revealed statistically significant genetic correlations among several traits, including CTMax and fork length, CTMax and DI, and family-level mean CTMax measured at 15 °C and 18 °C rearing temperatures. These correlations suggest the presence of shared genetic influences among traits. To test this hypothesis with genetic data, we evaluated overlap among genes identified in GWAS analyses for each trait and assessed SNP-level effect sizes for both traits using the PLACO+ framework. Despite the large number of genes linked to the top 300 SNPs per GWAS (90–144 genes), there was limited overlap among analyses: traits shared only 1–6 trait-associated loci (Table 1). However, given the large background set of possible genes (over 23,000) in LD with variant SNPs, several comparisons of overlap exceeded expectations compared to the null expectation of random association. These included overlaps that matched pedigree-based correlations, such as between the genetic-only associations for CTMax and: (1) its GxE associations for CTMax, (2) genetic-only associations for fork length, and (3) low-high progenitor F_ST_ divergence. Additionally, GxE associations for CTMax overlapped significantly with GxE associations for fork length. In contrast, SNP-level comparisons using PLACO+ found no SNPs that passed genome-wide nor chromosomal level thresholds between any comparisons (Fig. S6-11).

**Table 1.**
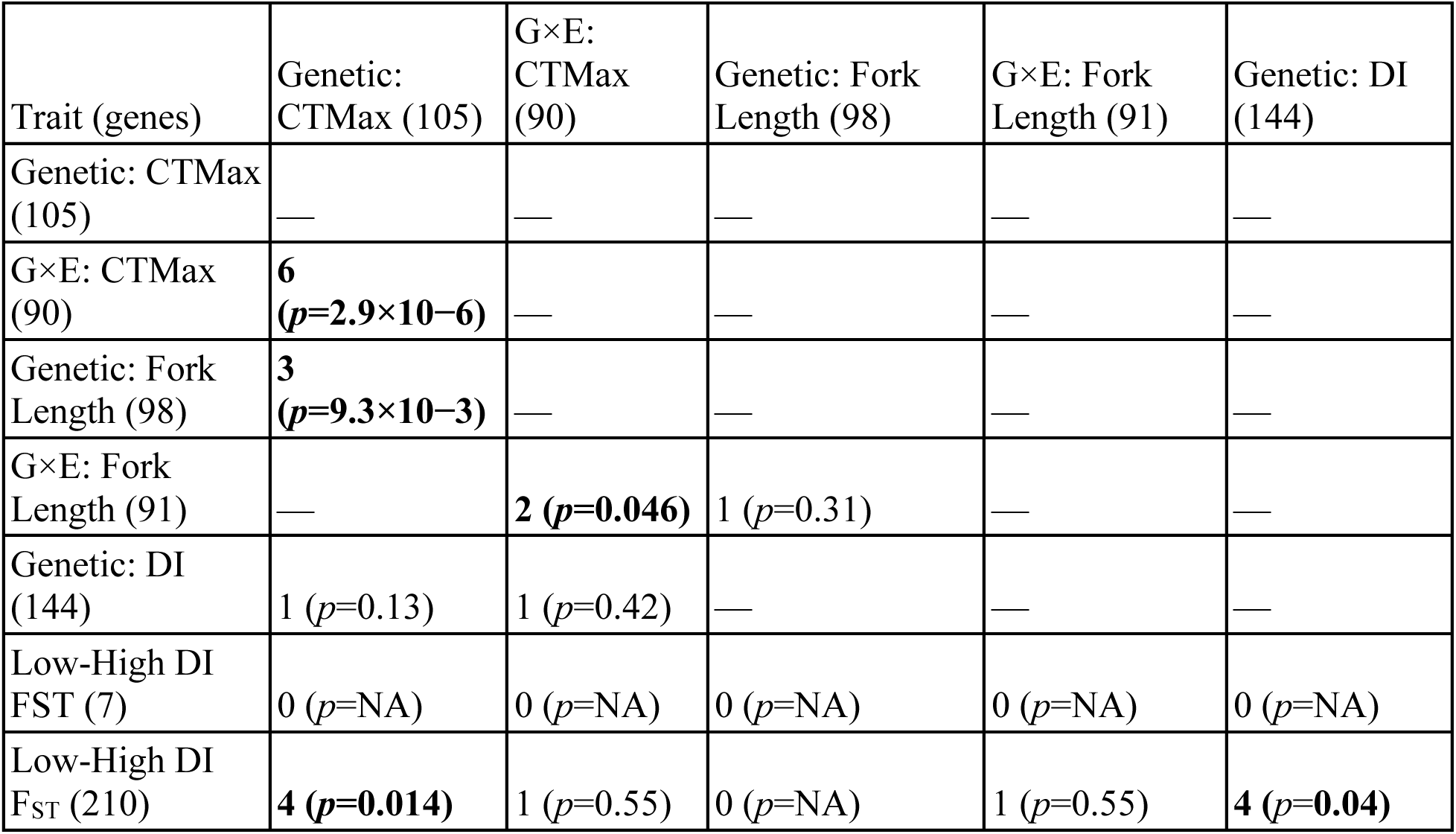
Pairwise overlap of genes associated with each trait in GWAS analyses, including for CTMax, fork length, and domestication index (DI). Numbers in parentheses indicate the total number of genes that were in LD with the top 300 SNPs identified for each GWAS. For low-high progenitor F_ST_, we compared overlap of 300 SNPs (7 genes) from the top windows, as well as SNPs above the 0.1 F_ST_ threshold (210 genes). The total number of overlapping genes between trait pairs is reported, with associated hypergeometric test p-values.

### Overlap of GWAS candidate genes with transcriptomic and epigenomic responses

To further explore genes associated with genetic and GxE effects on CTMax and DI, we tested whether candidate genes from GWAS analyses were significantly enriched for genes that were found in Griffiths et al., (2026) to be differentially expressed or differentially methylated between DI groups or between groups reared at different temperatures. We found that most GWAS gene sets were not significantly enriched for genes that were differentially expressed or differentially methylated between DI groups or between rearing temperatures (Table 2 & 3). However, the genes with genetic (GWAS) associations with DI were significantly enriched for genes that were differentially expressed between DI groups (18 genes shared between the two groups, which is more than expected by chance p=0.001).

**Table 2.** Pairwise overlap of genes associated with variation in CTMax identified from GWAS analyses and genes differentially expressed (DEGs) or differentially methylated (DMRs) in response to rearing temperature (15 ℃ vs. 18 ℃) or to rearing temperature × domestication index (DI) interactions. Numbers in parentheses in the row and column headers indicate the total number of candidate genes from GWAS or the total number of DEGs/DMRs from Griffiths et al., (2026). GWAS candidate genes from genetic associations with CTMax were defined using a −log₁₀(p) ≥ 3.5 threshold, whereas candidate genes from GxE associations with CTMax were defined using a chromosome-level significance threshold. For each comparison, the number of overlapping genes is shown, with the associated hypergeometric p-value provided in parentheses.

|  | Genetic: CTMax (209) | GxE: CTMax (247) |
| --- | --- | --- |
| Differentially expressed between rearing temperatures (1602) | 12 ($p=1$) | 15 ($p=1$) |
| Differentially expressed Temperature-by-DI interaction (26) | 1 ($p=0.48$) | 1 ($p=0.54$) |
| Differentially methylated between rearing temperatures (1244) | 22 ($p=1$) | 11 ($p=1$) |

**Table 3.** Pairwise overlap of genes associated with domestication index (DI) identified from GWAS analyses or progenitor F_ST_ and genes differentially expressed (DEGs) or differentially methylated (DMRs) between low or high DI fish or to rearing temperature × domestication index (DI) interactions. Numbers in parentheses in the row and column headers indicate the total number of candidate genes from GWAS or the total number of DEGs/DMRs from Griffiths et al., (2026). GWAS candidate genes from the genetic association with DI were defined using a using a chromosome-level significance threshold, whereas candidate genes for F_ST_ was based on values above 0.1. For each comparison, the number of overlapping genes is shown, with the associated hypergeometric p-value provided in parentheses.

| | Genetic: DI (192) | Low-High DI $F_{ST}$ (210) |
| --- | --- | --- |
| Differentially expressed between DI groups (353) | **18 ($p=1.1 \times 10^{-3}$)** | 11 ($p=0.27$) |
| Differentially expressed Temperature-by-DI interaction (26) | 1 ($p=0.45$) | 0 |
| Differentially methylated between DI groups (965) | 14 ($p=1$) | 25 ($p=1$) |

## Discussion

As captive-reared individuals are released into the wild they may face altered environments or a legacy of domestication selection that may cause traits to be mismatched to contemporary conditions. In the Delta Smelt hatchery, recent cohorts have relied on captive-born broodstock due to the scarcity of wild individuals and captive rearing temperatures are stable and cool during the summer. However, upon release, captive Delta Smelt will be introduced into a habitat that has been rapidly warming, where critical thermal thresholds dictate survival (Davis et al., 2024). Here, we assessed whether captive Delta Smelt harbor the genetic variation necessary to adapt to elevated temperatures, and how rearing temperature, body size, and domestication index influence this genetic infrastructure. We discovered some thermal acclimation ability (plasticity) during early life rearing at elevated temperature, but this was accompanied by a reduced plastic response in more highly domesticated Delta Smelt. Importantly, our companion study (Griffiths et al., 2026) found that both low and high DI fish still share similar molecular responses (transcriptomic and methylomic) to rearing temperatures, suggesting that the genomic mechanisms regulating thermal acclimation have not diverged. Genetic variation for thermal tolerance was reduced under warmer temperatures, signaling a constraint on adaptation potential. While few loci significantly contributed to the genomic architecture of thermal tolerance across both rearing temperatures, we identified extensive loci with significant rearing temperature-dependent effects, indicating that the genomic architecture is instead highly contingent on acclimation conditions. Furthermore, we demonstrate the influence of domestication selection; between low and high DI fish we detected loci with large allele frequency differences, as well as divergence in their transcriptomes and methylomes (Griffiths et al. 2026). However, we found minimal overlap between loci associated with domestication and thermal tolerance, suggesting that these two traits possess separate genetic underpinnings. We summarize these findings, coupled with those reported in the companion study Griffiths et al., (2026) in a conceptual model (Fig. 9).

**Figure 9:**
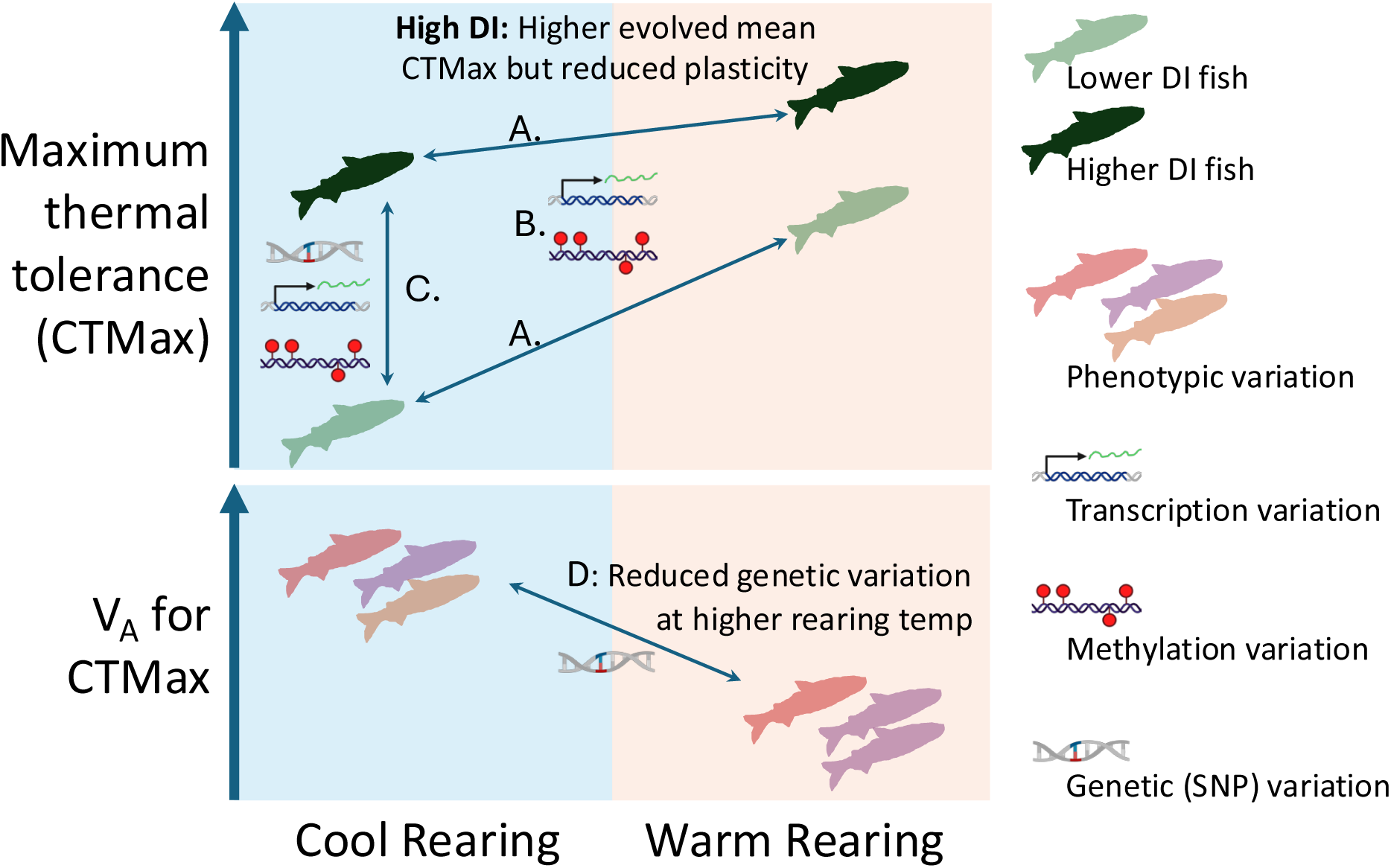
Conceptual summary of main findings from this study and Griffiths et al., (2026). A) Delta Smelt can increase their thermal tolerance (CTMax) if reared at elevated temperature (acclimation or phenotypic plasticity). While higher domestication index (DI; black) fish have higher mean thermal tolerance at both rearing temperatures, they have reduced thermal plasticity compared to lower DI (green) fish (slope connecting lower DI fish between cool and warm rearing is steeper than for the slope connecting higher DI fish). B) Rearing at different temperatures induced transcriptomic and methylomic changes but with similar responses among low and high DI fish (see Griffiths et al., 2026). C) Phenotypic variation in CTMax that distinguishes low from high DI fish is likely underpinned by evolved genetic variation that manifest as transcriptomic and methylomic variation. D) Additive genetic variation in thermal resilience is reduced in warmer rearing environments.

### Drivers of Thermal Tolerance and Adaptive Potential in Delta Smelt

The ability for fish to increase their upper temperature tolerance after acclimation to elevated temperatures is an important physiological mechanism for maintaining fitness, such as during heat waves in the summer. Consistent with previous findings (Griffiths et al., 2026), Delta Smelt showed a modest yet statistically significant capacity for thermal plasticity (acclimation), where fish reared at 18°C exhibited an increase in CTMax (0.6°C) compared to those reared at 15°C (Fig. 1). The upper limit of CTMax that we observed in this study was around 30-31°C, which was consistent across both rearing temperatures and previous studies (Komoroske et al., 2014). This temperature may therefore represent a physiological barrier for the species (Pörtner et al., 2017). Although this upper thermal limit (30-31°C) exceeds temperatures that caused total mortality during experimental releases (26°C), it is widely acknowledged that CTMax methodology does not accurately reflect sustained exposure to extreme heat (Bartlett et al., 2022; Brauner & Richards, 2020).

Domestication appears to have altered thermal physiology within Delta Smelt, insofar as more highly domesticated fish (high DI) exhibited higher CTMax than lower DI fish at both rearing temperatures. There was a 0.24°C increase in CTMax with each 1-unit increase of DI (each DI unit represents 1 generation spent in the hatchery). This suggests that genetic variation for thermal tolerance may be partitioning among fish with different degrees of hatchery ancestry. However, Delta Smelt are reared at constant temperatures that are lower than temperatures in the wild, such that directional selection for increased thermal tolerance seems unlikely. Instead, thermal tolerance may be genetically correlated to other traits under selection in the hatchery (e.g., metabolic rates, feeding capacity, stress tolerance).

As the currency of evolutionary change, genetic diversity is important for population persistence during adjustment to environmental shifts; consequently, a loss of variation - particularly at quantitative trait loci - can compromise evolutionary potential. To investigate standing genetic variation for thermal tolerance for captive-reared Delta Smelt, we used both pedigree- and genomics-based estimates. We observed heritabilities within the same range for both estimates (0.15 and 0.26 pedigree-based, and 0.14 genomics based). This suggests that while variation in thermal tolerance has a large environmentally-dependent component, there is a significant but modest genetic basis that can be acted upon by selection. The heritability of CTMax measured in Delta Smelt is lower than that estimated in other species such as Atlantic salmon (0.47) and rainbow trout (0.41) (Benfey et al., 2024; Perry et al., 2005).

Despite similar heritabilities for thermal tolerance between fish reared at the two rearing temperatures, additive genetic variation and evolvability were reduced at the warmer temperature (18 °C). Phenotypic variation was also reduced in the warmer temperature (Fig. 1A), corresponding with reduced residual and environmental variation in the animal model (Table S1). This reduction in non-additive genetic variance likely contributed to the similar heritability estimates between the two temperatures. Since the evolutionary response to selection depends on the amount of additive genetic variation, adaptation for higher thermal tolerance will be more limited at the warmer rearing temperature. If CTMax was under continuous directional selection, our estimates of evolvability correspond to a 0.25 % and 0.06 % change in CTMax per generation, at rearing temperatures 15°C and 18°C, respectively. This suggests that while Delta Smelt possesses standing genetic variation to modify thermal tolerance, this capacity will be constrained under warmer conditions.

We detected evidence for genetic variation for thermal tolerance plasticity (genetic associations with CTMax that varied depending on rearing temperature), suggesting genotypes differ in their capacity to acclimate to warmer environments. Our previous work demonstrated that Delta Smelt that were more highly domesticated had reduced capacity for thermal acclimation (reduced phenotypic plasticity) compared to less domesticated fish (Griffiths et al., 2026). Stable rearing temperatures in the Delta Smelt hatchery may have, over generations, contributed to a loss of thermal plasticity in more highly domesticated fish. Despite this, we were unable to detect differences in additive genetic variation and heritability for CTMax between low, medium, and high DI fish. Although higher domesticated fish may have lost genetic variation for thermal plasticity, they still maintain the ability to evolve a higher mean thermal tolerance.

### Genomic architecture of thermal tolerance

Given evidence of thermal tolerance heritability in Delta Smelt, we used GWAS to reveal the number and identity of associated genetic loci. We found only a few SNPs that passed statistically significance thresholds for association with thermal tolerance across both rearing temperatures, suggesting variance in this trait is most likely polygenic – including many loci of small effect. Identifying the specific loci that could be targets of selection for thermal tolerance in fish remains a challenging task (Gonen et al., 2024; Ignatz et al., 2025). Heat tolerance has been demonstrated to be a highly polygenic trait in other species (Fuller et al., 2020; Gonen et al., 2024; Griffiths et al., 2020; Michalak et al., 2019; Tobler et al., 2014). SNPs associated with polygenic traits are difficult to detect, since the effect size of each contributing locus is small (Yeaman, 2015). Since the trait is likely polygenic, we sought to test whether a wider set of SNPs with individually lower association values might also implicate mechanisms underlying variation in thermal physiology. Indeed, tests for functional enrichment of the top 0.04% of the SNPs identified many more functions and cellular components, which were consistent with other studies that have shown that physiological and cellular heat stress has been characterized by loss of ionic and osmotic balance (reviewed by Evans, 2008) and the compromise of lipid membrane fluidity (Hochachka & Somero, 2002).

While we detected a relatively limited number of loci with consistent effects on CTMax across rearing temperatures, we identified a large number of loci where the genetic association with CTMax depended on the rearing temperature. This environmental dependence (rearing temperature) of genotype on phenotype is consistent with our finding that additive genetic variation for CTMax differed between the two rearing temperatures. These temperature-dependent loci were enriched for a wide-range of functional categories including gene regulation and structure and cellular organization (i.e., chromatin organization, protein-DNA organization, and cellular component organization), suggesting that loci with a GxE effect may involve structural changes to the genome and rearrangement of cellular structures. These loci may also cause adjustments to metabolic processes (i.e., ATP-dependent activity) and cell signaling and communication (i.e., cell-cell signaling, endopeptidase inhibitor activity, and enzyme regulator activity).

In a companion study, we demonstrated that elevated rearing temperatures resulted in large transcriptomic and methylomic changes in juvenile Delta Smelt (Griffiths et al., 2026). However, SNPs associated with CTMax and with GxE CTMax were not linked to the same genes as those involved in temperature-induced transcriptional and epigenetic changes.

Therefore, the segregating genetic variation associated with thermal tolerance limits (i.e., CTMax) is in different molecular machinery than that governing physiological acclimation during early life development. While rearing at 18°C generally primed fish to develop higher CTMax values, the GWAS and DEG/DMR methodology were measuring two related, but slightly different traits: acute upper thermal tolerance limits (CTMax) and long-term acclimation to elevated temperatures.

### Fork length, covariates, and genomic architecture

Fork length in Delta Smelt is a complex trait influenced by both rearing temperature and genetic variation. Fork length showed a positive plastic response to rearing temperature, where larger fish emerged at higher rearing temperature (Fig. 1C). Warm rearing temperatures often increases metabolic rates, which can lead to faster growth (Wood & McDonald, 1997). However, we also estimated large pedigree- and genomic-based heritabilities for this trait (0.58 and 0.50 pedigree-based, 0.30 genomic-based), consistent with genetic variation contributing to trait variation. Indeed, we detected one large-effect locus that overlapped with four genes, two of which are annotated, but with functions not typically associated with size or growth. An additional signal of subtle polygenic variation associated with this trait was distributed across multiple chromosomes. Functional enrichment of genes in these peaks suggests that cellular structure and organization, metabolism, homeostasis, and cell signaling contribute to variation in fish length. Compared to the number of loci associated with genetic variation for fork length, there were many more loci that had a rearing temperature dependent effect on fork length. These GxE loci for fork length were enriched for cellular structure and metabolism functions, as well as protein binding and regulation (such as GTPase binding, which acts as a molecular switch known to control a variety of functions, including growth; (Satoh, 2020)).

Fork length displayed strong negative phenotypic and genetic correlations with CTMax, indicating that larger Delta Smelt have reduced thermal tolerance. Similar findings have also been found in other salmon species (Bartlett et al., 2022; Benfey et al., 2024; Debes et al., 2021). Our observed negative genetic correlation between these two traits suggests a potential shared genetic basis, with the implication that selection may be constrained from simultaneously increasing both fork length and CTMax. Although this trade-off could be a concern for fish in a warming climate, we found that this effect was diminished at warmer rearing temperatures. This may be a result of reduced genetic variation for CTMax at 18 °C, and phenotypic plasticity may play a larger role in determining the final phenotype. At the genomic level, we found some significant overlap in genes for CTMax and fork length GWAS (Table 1), which is consistent with at least some shared genetic basis.

### Signatures of domestication selection

Despite careful genetic management, evidence for domestication in the Delta Smelt hatchery has been observed for a variety of traits, such as reproductive success, growth, and survival. In the hatchery, Delta Smelt spawners that have greater hatchery ancestry tend to produce more offspring that survive to reproductive age compared to fish with shallower hatchery ancestry (Finger et al., 2018; LaCava et al., 2023). However, in this study, we found that larval (71-79 dpf) survival rates among families were similar across rearing temperatures and DI categories, and that DI had only a small effect on size. In contrast, previous work found pronounced differences in size and survival among low and high DI fish (Chase et al., 2024; Ellison et al., 2023), but those studies included very low DI fish that were not available for our experiments.

We identified genomic divergence within both the progenitor and offspring populations. We detected elevated F_ST_ differentiation between low and high DI progenitors as well as SNPs associated with continuous variation in DI within the offspring population. These results may represent an underlying genomic basis for divergence in traits within hatchery populations (Finger et al., 2018; Griffiths et al., 2026; LaCava et al., 2023). Divergent loci were functionally enriched in categories related to development and morphogenesis (i.e., multicellular organism development, anatomical structural development, and developmental processes), which suggests selection on development and growth traits. These loci may contribute to the divergence in physiology and morphology that we observed between low and high DI fish, but may also influence other morphological or developmental traits that we did not measure (Fig. 1C). For example, loci distinguishing low from high DI fish were linked to genes that were functionally enriched for GO categories related to synaptic and neural signaling (i.e., synaptic signaling, cell-cell signaling, and chemical synaptic transmission), which suggests that domestication has altered neurological and behavioral systems as observed in other domesticated fish (e.g., (Ghazal et al., 2025)).

Genomic variation distinguishing low from high DI fish may be associated with divergence in gene expression between low and high DI fish. We detected significant overlap in the gene set identified by GWAS association with DI and the gene set that was differentially expressed between low and high domesticated fish from Griffiths et al. (2026). Our results are consistent with domestication selection being a complex process that influences both genetic variation and molecular response pathways. Accordingly, one may consider the influence of hatchery conditions not only on survival but also as a filter for genotypic variation and as a lever that influences the evolutionary forces that govern the fate of that variation.

### Conclusion and Hatchery Implications

Elevated and sustained resilience to environmental variation should be a goal of most conservation hatcheries; especially those that hope to release captive animals back into environments being altered by climate change (Bradford et al., 2025; Fraser, 2008). Maintaining or enriching for thermal tolerance in Delta Smelt will be critical for the success of the Delta Smelt supplementation program. This is especially apparent given that recent experimental releases have experienced high mortality driven by temperature extremes (Baerwald et al., 2023; Davis et al., 2024; U.S. Fish and Wildlife Service, 2020). Our results indicate the existence of standing genetic variation which could be the substrate for natural selection to adaptively increase thermal tolerance. Unfortunately, this adaptive capacity may be constrained under warmer conditions. Delta Smelt also possess the capacity for improving thermal tolerance through acclimation, and the presence of many loci with rearing temperature-dependent impacts on thermal tolerance suggests there is genetic variation for thermal plasticity. Thermal performance traits in Delta Smelt are therefore a combination of genetic variation, genotype by environment interactions, and acclimation, making it challenging to predict the interventions that will maximize the success of supplementation. Additionally, simple marker-assisted breeding will be challenging, given the highly polygenic nature of thermal tolerance and the potential for complex interactions with other genetic variation important for fitness in the wild. Given that all Delta Smelt now originate from the hatchery, future research should track phenotypic traits and domestication indices following supplemental releases to determine how these factors influence fitness in the wild.

## Data Accessibility

Sequencing data are deposited in the National Center for Biotechnology Information’s Short Reads Archive (BioProject PRJNA1152625). Scripts and code for physiological and genomic analyses are available on GitHub: https://github.com/JoannaGriffiths/DeltaSmelt_Genetics. The progenitor reference panel can be found on Dryad (DOI: 10.5061/dryad.qz612jmx7).

## Acknowledgements

This research was supported by the California Department of Fish and Wildlife (Prop 1: Watershed Restoration Grant Program / Delta Water Quality and Ecosystem Restoration Program. Grant Q1996041, PIs Dr. Andrew Whitehead, Dr. Nann Fangue, Dr. Tien-Chieh Hung). Additional funding provided by the UC Agricultural Experiment Station (2098- H) to NAF. We thank current and former FCCL staff for fish maintenance (especially Luke Ellison and William Mulvaney). We thank UC Davis staff that helped with physiological measurements (Jenn Roach, Dr. Nicole McNabb, Ashley De La Torre, and Anthony Tercero). The sequencing was carried out at the UC Davis Genome Center DNA Technologies and Expression Analysis Core, supported by NIH Shared Instrumentation Grant 1S10OD010786-01. We acknowledge the High Performance Computing Core Facility at the University of California, Davis, for providing computational resources that have contributed to the research results reported in this paper.

